# The evolution of plant actins

**DOI:** 10.64898/2026.04.22.720131

**Authors:** J. Lamers, D. Pereira, M. Kaper, J. Sprakel

## Abstract

Why plants have evolved large families of highly similar actin isoforms, and what functional distinctions sustain their retention, remains an unresolved problem in cytoskeletal biology. Actin is among the most ancient and conserved proteins in eukaryotes, yet plant genomes encode an unusually high number of closely related paralogs whose specialization is poorly understood. Here, we reconstruct the evolutionary history of plant actins across the green lineage by combining sequence retrieval, phylogenetics, comparative sequence analysis, and structural mapping. We find that plant actin diversification arose multiple times and, strikingly, relatively late in the history of this ancient protein family. A major duplication produced two deeply conserved seed-plant actin clades, corresponding to the previously described vegetative and reproductive actins; here, we refer to these as Type I and Type II to reflect their phylogenetic relationships. Across the green lineage, increases in actin copy number track key transitions in plant complexity, particularly in tracheophytes. Despite this diversification, only a limited set of conserved substitutions distinguishes major actin lineages. Mapping sequence variation onto actin structures reveals a prominent surface patch that appears permissive to mutation, suggesting relaxed functional constraint, whereas isoform-specific changes cluster at sites likely to influence filament stability, turnover, and treadmilling. A finer-scale analysis in Brassicales pinpoints recent substitutions predicted to alter hydrogen-bonding patterns within monomeric and filamentous actin. Together, our results argue that plant actin diversification is not redundant drift, but a recurrent evolutionary strategy that preserves the actin core while tuning the intrinsic biophysical behavior of f-actin.

## Introduction

Actin is one of the most abundant and highly conserved eukaryotic signature proteins. The discovery of actin orthologs in Asgard archaea^1^ points to an ancient origin, yet actin sequences have remained strikingly conserved across the eukaryotic domain despite this deep evolutionary history. The plant lineage exhibits a striking expansion of actin isoform diversity, often encoding larger, more diverse actin gene families than those of animals and fungi^2,3^. This is particularly astute in seed plants, which have undergone repeated genome duplications, with several species expressing 10-20 bona fide actin isoforms^4^. Most plant genomes across the green lineage encode numerous actin paralogs, often differing by only a few percent at the amino acid level^4^. This combination of extreme sequence similarity and substantial gene family expansion presents a yet unresolved puzzle: why has actin diversified so extensively in plants, and what functional distinctions justify the maintenance of so many subtly different isoforms?

In animals, different skeletal^5–8^ or cytosolic^9,10^, actin isoforms are often broadly interchangeable within their respective group. Functional differences arising primarily from differential expression patterns rather than intrinsic biochemical specialization. By contrast, in plants actin isoforms appear to be non-redundant^11,12^. Specific isoforms are required for distinct developmental contexts, including tip growth, cell elongation, and reproductive processes^13–15^. The developmental phenotypes resulting from the deletion of one actin isoform cannot be rescued by complementation with another isoform; rather, they can lead to much more severe defects^11,12^. This suggests that, even though sequence differences between plant actin isoforms within the same species are small, they encode functionally meaningful distinctions. Yet the mechanistic basis of how and where this isoactin specialization is encoded in the protein sequence or structure remains poorly understood.

We identify two main, non-mutually exclusive explanations for plant actin diversification. First, subtle changes in sequence between isoactins may lead to the specialization of the actin interactome. Actin-binding proteins (ABPs), which regulate filament nucleation, elongation, severing, bundling, and spatial organization, could exhibit differential affinities for distinct actin isoforms, thereby creating specialized cytoskeletal networks tailored to particular cellular contexts. Second, subtle sequence variation may directly tune the intrinsic biophysical properties of f-actin, including polymerization kinetics, filament turnover, and higher-order organization. Even small changes at key residues can alter filament dynamics, which are amplified at the network level, producing emergent functional differences needed for distinct cellular requirements.

Both possible explanations, by consequence, predict that actin isoforms co-evolved with the complexity of the plant. As plants evolved more complex body plans, with more specialized cell types and organs, this could have led to the need to evolve increasing numbers of tailored isoforms. Even though the last thorough exploration of plant actin evolution was published before the first whole plant genome was available^16^, these studies provided clues that support this notion. In particular, flowering plant actins were found to occupy two main clades, whose expression patterns correlated with either vegetative or reproductive tissues. This suggested that, as plants evolved more complex reproductive strategies, actins diversified in tandem.

The rapid accumulation of high-quality plant genomes across the green lineage now presents an opportunity to address these hypotheses from a more comprehensive evolutionary perspective. Across the green lineage, actin gene families have undergone lineage-specific expansions, contractions, and diversifications. This phylogenetic breadth enables the identification of conserved and lineage-specific sequence features, as well as the trajectories of molecular evolution. Importantly, patterns of sequence conservation and divergence can reveal residues under selective constraint, sites of adaptive change, and co-evolving positions that may mediate isoform-specific functions.

Here, we exploit the rapidly expanding genomic resources to systematically analyze the evolution of plant actin isoforms. By integrating phylogenetics, comparative sequence analysis, and structural mapping, we identify patterns of amino acid variations across the green lineage and evaluate their potential functional significance. Through this data-driven approach, we aim to generate testable hypotheses linking sequence variation to functional specialization. Our results suggest that plant actin diversification is neither neutral nor redundant but instead reflects a finely tuned balance between conservation of core function and adaptive divergence. We find that actin diversification occurred several times in the evolution of the green lineage and correlates with important evolutionary transitions in plant anatomy and habit. Remarkably, our analysis suggests that plant isoactin diversification occurred relatively recently as compared to the entire evolutionary history of the ancient actin protein family. By structural mapping of mutational frequencies, we identify a patch on the three-dimensional actin fold that, unlike all other parts of the actin structure, appears to have been free from the strict evolutionary pressure that shapes the overall evolutionary trajectory of actins. This indicates the presence of an actin site at which mutations or interventions are relatively permissible. Moreover, we find that more subtle mutational hotspots that separate distinct actin isoforms are more likely to be involved in the intrinsic f-actin biophysics, such as filament stability and turnover, rather than in shaping the actin interactome. Our work provides a comprehensive picture of plant actin evolution based on currently available genomes and offers tangible leads for dissecting the molecular mechanisms underlying the functional diversification of plant isoactins.

## Results

### Methodology

Despite being a key actor in many cellular processes across the green lineage, to the best of our understanding, the latest comprehensive exploration of the evolution of plant actins stems from 1999^17^. At this time, no whole plant genomes were available, the first being published in 2000^18^, and only a few (partial) actins were known. Moreover, structural and computational biology were in its early stages, and the Asgard archaea and their actins were not yet discovered. Motivated by the surge in the availability of whole-plant genomes in the past decade, we decided to revisit the evolutionary history of plant actins. We searched for actins in existing proteomes of 5 Asgard archaea, 2 opisthokonts, 8 red algae (rhodophyta) and 165 plant species of major taxonomic clades (Supp. Table 1). These proteomes were carefully selected for being of high quality, and numerically balanced among the plant lineages (see Experimental methods). Obtaining a clean, robust, and comprehensive phylogeny of actin isoforms in the plant lineage is challenging for several reasons. First, actins and Actin Related Proteins (ARPs), in which actin is the major domain, are very similar at a sequence level and are not always easily distinguished. Secondly, actin is considered one of the most sequence conserved eukaryotic proteins; for example, animal and plant actins show only approximately 10% sequence divergences, and differences within the plant lineage are much smaller. The high level of evolutionary conservation makes the phylogeny very sensitive to small sequencing and annotation errors. Actins with sequence errors are clustered together as an outgroup. We tackled these challenges by creating a Hidden Markov Model (HMM) profile using five different actins among the tree of life (plants, metazoa, fungi, oomycetes) to extract actin-like proteins from proteomes. Next, we filtered the hits using a minimal threshold (full sequence score >70). This threshold was arbitrarily chosen, but amply included reported pseudoactins (e.g. AtACT9 = 640) and highly divergent actins in Chlamydomonales (e.g. jgi|Volafr1|993|GIL43051.1 = 587). We performed HMM searches on these filtered hits with multiple existing PANTHER profiles (ACT, ARP1/2/3/4/6/9, ACTL7 and centractin) and used the HMM scores for principal component analysis (PCA, Supp. Fig. 1A,B, Supp. Table 4). Based on the known *Arabidopsis thaliana* proteins, their strongly conserved length (∼375-377 amino acids) and their conserved C-terminus (HRKCF), we identified PCA group 6 as genuine actin proteins (Supp. Fig. 1C). The proteins in this group were all genuine actins but still contained proteins with annotation errors. In particular, we note that the first exon was often missed and the last exon often partially annotated as an intron. Preferably, those sequences were curated by BLAST, but 91 sequences required manual curation against genomic data (Supp. Table 2). 42 actins were eventually removed as neither BLAST or manual curation led to complete actin sequences (Supp Table 3). As few annotated lycophytes genomes are available, we made an extra effort to include as many actins as possible. Therefore, we manually identified actins using the conserved C-terminus in *Lycopodium clavatum* and *Isoetes taiwanensis*. Proteins with exact sequence duplications within the same organisms were removed. To the best of our knowledge, the strategy of combining HMM scores with PCA to select candidates has not been reported. Here, we show that this is a very powerful approach to find protein sequences with relaxed thresholds. This led to the identification of protein sequences that were severely truncated/misannotated and would have been missed with more stringent or arbitrary thresholds. The subsequent PCA using multiple HMM profiles then allows proper separation of actins and ARPs.

### Phylogeny

Next, protein sequences were aligned using ClustalOmega and the phylogenetic tree was created with the iqTree2 software (Supp. Table 5). An interactive version of the phylogeny is available online (https://itol.embl.de/shared/JasperLamers). The tree was rooted on an Asgard archaea actin (2951803_UPI0027A42375), as these form the outgroup to all Eukaryotes. Early studies of *Arabidopsis* actins revealed two types of actin, which, based on their spatial expression patterns, were termed vegetative actins (ACT2, 7 & 8) and reproductive actins (ACT1, 3, 4, 11, and 12)^19^. Strikingly, this division also emerges from the phylogeny of protein sequences and is not limited to *Arabidopsis* but found in all angiosperms^17^. The correlation between expression patterns and protein sequence, suggests that these actin types may have distinct functions. In our tree, we also identify two distinct classes (99% bootstrap support, Fig. 1A). Interestingly, we also find the reproductive-type actins in gymnosperms. This reveals that this diversification in actins is a seed-plant specific adaptation, rather than an angiosperm-specific adaptation as previously thought. The reproductive actins of gymnosperms cluster together and form an outgroup to the reproductive actins of angiosperms. This matches the evolutionary trajectories of these lineages. This finding aligns well with the earlier estimation that vegetative and reproductive actins split ∼400 million years ago^20^. This moment coincides with the emergence of the earliest seed plants ∼385-365 million years ago, while flowering plants only emerged 150 million years ago. Taken together, we argue that the identified gymnosperm actins are *bona fide* reproductive-type actins.

**Figure 1.**
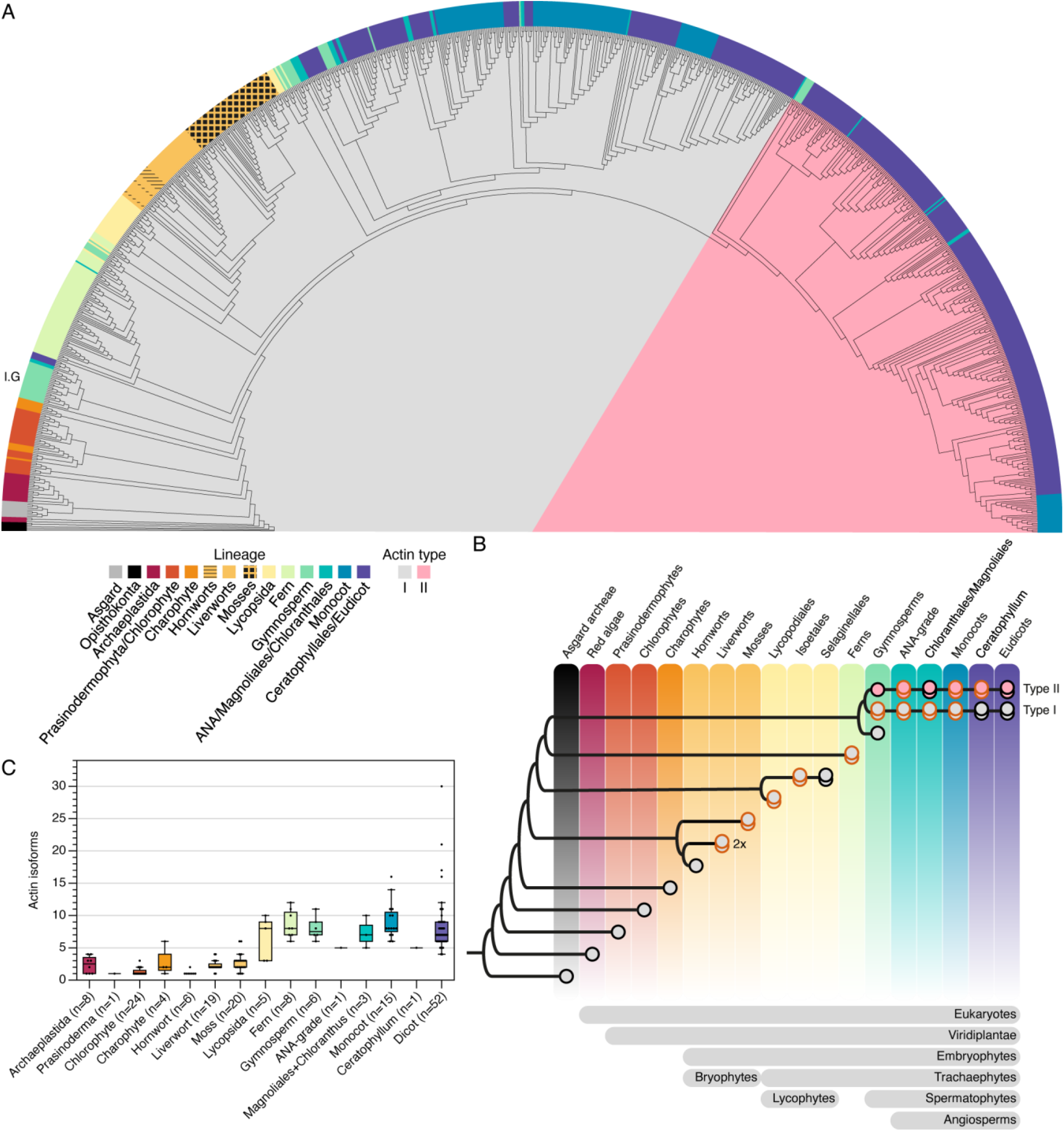
Ancestors of most plant lineages possessed only a single actin gene. (A) A phylogenetic tree covering 904 actins from 173 plant species, 2 opisthokonts and 5 asgard archaea. (B) A summary tree of actin evolution. Multiple dots indicate the existence of at least two clades within an organism. Dots with red borders indicate *de novo* multiplication events, in which the ancestors most likely had a single actin copy of that type. The multiplier shows that *de novo* multiplication events happened multiple times in the lineage. Gray and pink dots indicate, type I and II actins, respectively. (C) The unique actin protein copy number per selected plant lineage.

This also implies that all lineages that split before the gymnosperms do not contain reproductive actins, while these plants clearly do reproduce. This indicates that the terminology is seed plant-centric, or initially Angiosperm-centric, and excludes many plants lineages. Moreover, this nomenclature was derived based on expression patterns, while it is now clear that vegetative actins are expressed in reproductive tissues and *vice versa*^19,21,22^. We thus argue that this terminology is outdated as it is confusing, especially for those who entered the field more recently. Hence, we propose that these actin types are referred to based on their phylogenetic grouping as Type I (formerly: vegetative) and Type II (formerly: reproductive) actins.

Strikingly, within the Type I actins, which represent the ancestral actin lineage, actin isoforms are strongly clustered per plant lineage with, starting (on the left), segmented clusters containing actins belonging to the (1) red algae/opisthokonts, (2) chlorophytes/charophytes, (3) gymnosperms (I.G), (4) ferns, (5) lycophytes + bryophytes and (6) gymnosperms and (7) angiosperms. Gymnosperms are an interesting case as they are the only non-angiosperms with two distinct type I clusters. Cluster I.G forms an outgroup to all embryophytes and contains gymnosperms, and few (eu)dicots (97% bootstrap support). The (eu)dicot actins in this cluster are not found in closely related species and we hypothesized these to be sequencing errors or pseudogenes. We tested this hypothesis by first generating AlphaFold3 models^23^ of a functional Arabidopsis actin gene (AtACT7), a hypothesized Arabidopsis pseudogene (AtACT9), a gymnosperm and eudicot actin of cluster I.G. At a first glance, AtACT9 still resembles an actin protein, but it has regions that are strongly tilted versus the AtACT7 (Supp. Fig. 2A, RMSD = 5.7 Å). No large differences were observed between the gymnosperm actin and AtACT7 (RMSD = 0.3 Å), but the dicot I.G actin also showed deviations (RMSD = 1.8 Å). This is the first indication that the latter are likely pseudogenes. Next, we compared the sequences of I.G actins to all other included plant actins in this study. We first plotted the AA frequencies of four I.G actins (Supp. Fig. 2B). Next, we created a boxplot of these frequencies for all actins in cluster I.G (Supp. Fig. 2C). Both analyses showed that the dicot actins in cluster I.G contain many rare AAs, a pattern that resembles the pseudogene AtACT9. On the other hand, the gymnosperm I.G actins contain some rare AAs, but not as many as the pseudogenes. This sequence analysis is thus in line with the structural predictions and the evolutionary conservation and we argue that these are genuine gymnosperm-specific, and likely functional, actins; the function of this gymnosperm specific adaptation is unknown. Next, we reconstituted how the actin duplication and diversification led to the diversity of distinct actin isoforms (Fig. 1B). The clear segmentation between plant lineages, implies that nearly every non-angiosperm plant lineage originated from an organism with only one actin isoform that has been conserved throughout evolution. Such a strong segregation has also been observed in opisthokont actin evolution, in which mammals, insects, fungi, nematodes and echinodermata form independent subclusters^24,25^. We have identified at least 13 events in which such ancient organisms evolved into extant organisms with at least 2 actin isoforms. We counted actin types I and II separately and included only clades that contained multiple species. Therefore, recent duplication events in which multiple actins of one organism form a subclade in the phylogeny were not included in this estimation. We note that it is possible that the ancestral organism contained more than one actin, but if all but one were lost early in the evolution of the plant lineage while the other actin diversified, it now appears that the ancestral organism only had a single actin copy.

These findings imply that the evolution of actin isoforms occurred numerous times in evolutionary history and that the diversity of actin isoforms we observe today emerged rather recently. This seems somewhat counterintuitive for such an ancient protein, but it underlines the enormous evolutionary pressure on conserving the structure of actin to preserve its essential functions, while only residues appear highly tailorable to suit the organism or specific task within an organism. Extreme examples of such adaptations are found in the chlamydomonadales chlorophyte algae, which contain two actins, of which one is a canonical actin with 90% homology to human ACTB^26,27^. The other actin is strongly divergent and only 60-69% homologous to the canonical actin. The divergent actin is important for flagella development and organization, and thus represents a specific adaptation to meet a specific cellular function.

We hypothesize that actins have diversified independently many times to tailor them to specific needs. This hypothesis implies that the actin copy number must be positively correlated to the complexity of the body plan. Hence, we plotted the number of unique actin protein sequences per lineage sorted by their evolutionary divergence (Fig. 1C). This shows that number of actin isoforms is roughly constant between red algae and mosses, but rapidly increased in the tracheophytes. These have more complex body plans and distinct cell types compared to the bryophytes. Interestingly, while the Type II actins developed with the emergence of seed plants, the number of actin isoforms did not increase in seed plants. This implies that the sequence diversity of actin isoforms of spore producing tracheophytes (lycophytes and ferns) is lower and might be more redundant compared to the seed plants. This is supported by our phylogenetic tree, in which fern and lycophyte actin isoforms form intra-species clusters with up to 5 isoforms (Supp. Fig. 3).

### Evolutionary patterns in actin mutations

A possible reason for the enormous diversification of actin isoforms in the plant lineage is that these were required to accommodate important evolutionary transitions in body plan complexity or organismal habit. To this end, we ask if specific mutational substitutions co-occur with key evolutionary transitions. To extract these conserved substitutions, we compared actin proteins from plant lineages present before the split of a new lineage with those from all lineages that emerged after that split. For example, to identify the conserved changes that occurred during the evolution of the lycophytes, actins in lineages that formed before the lycophytes (prasinodermophyta-bryophyte) were grouped and compared with those in lineages that formed after this formation (lycophytes-eudicot). We calculate the amino acid frequency per position and extracted the positions in which the frequency changed |>70%|. For example, at alignment position 278, Asn is dominant (74%) in prasinodermophyta-bryophytes, but is changed to a Gln in lycophytes-eudicots (100%). This algorithm thus identifies positions where the dominance of a particular amino acid is gained or lost, for changes that are then conserved in subsequent evolution. It should be noted that this approach identifies only strongly conserved substitutions, not lineage-specific substitutions.

Most differences were found between red algae and prasinodermophyta/chlorophytes (Supp. Fig. 4). This comparison highlights the variation in red algae actin, compared to a much more conserved state of plant actin. Strikingly, we find that at this evolutionary split, 1-2 amino acids were added to the N-terminus of plant actin (Residue 8-10). Like the red algae, typical opisthokont actin is also 375 amino acids compared to 377 in plants. This indicates that the actin length increased with the evolution of the first plants. N-terminal extensions also occurred in animal skeletal actins (376 AAs) and more extremely in the hornwort *Anthoceros punctatus* (384 AAs). We initially expected a sequencing error, but this extension is robustly identified in two independent accessions (Bonn and Oxford) and in *A. punctatu*^28^. The N-terminus is relatively variable and displayed on the surface, but the biological relevance for N-terminal elongation in plant actin remains unknown.

In the following, we only compare actins in the green lineage to allow for a cleaner analysis of conserved substitutions (Fig. 2). This shows that only 19 residues have showed conserved substitutions during evolution, of which the majority are found in lineages that emerged before the angiosperms. Interestingly, some positions in gymnosperm actins are relatively variable compared to other lineages and contain intermediates between ferns and angiosperms (position 23, 91, 174, 375). We also investigated the evolution of Type II actins that emerged in the seed plants. The development of this actin type came with unique changes (position 143 and 247) and some that were already present in either ferns or gymnosperms (residue 14, 23, 91, 174, 375). As with Type I actins, no major substitutions occurred after the evolution of angiosperms. Interestingly, Type II actins retain the more ancient Alanine at position 375, while this transitioned to a dominant Serine in Type I. Taken together, this analysis shows that all conserved substitutions in both actin types happened before the evolution of angiosperms. However, as even actin isoforms of *Arabidopsis thaliana* are poorly understood, the biological relevance of these conserved changes in evolution remain a mystery. The effect of such substitutions has to be studied in the corresponding plant lineage, as we hypothesize that they serve specific functions in their host organism; unfortunately, for many lineages there are no distinct model organisms that are genetically accessible. Without dedicated efforts to develop these tools, the function of these substitutions will remain unclear.

**Figure 2.**
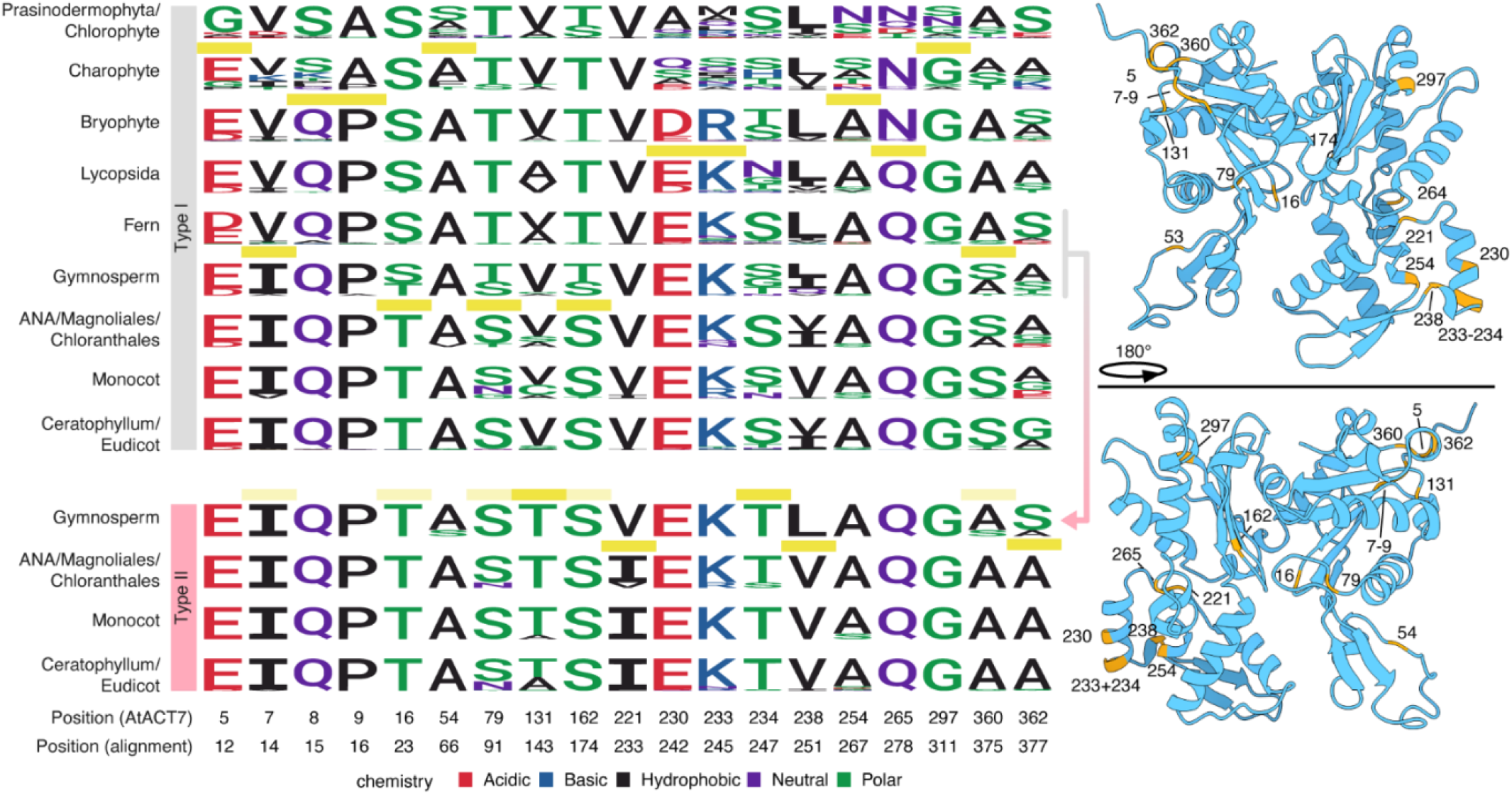
Conserved substitutions happened “early” in plant evolution. The major transitions in the actin sequences are mapped per major taxonomic clade and per actin type. Yellow bars indicate a transition of a specific amino acid. The opaque yellow bars indicate amino acids that were already (near)-dominant in ferns or gymnosperms and it remains uncertain whether these were substituted with the formation of type II actins or that these belonged to the ancient Type I actin that transitioned to a Type II actin. Gymnosperm cluster I.G was excluded from this comparison. The identified substitutions were mapped on the Arabidopsis ACT7 g-actin AlphaFold3 structure. Numbers indicate the alignment residue numbers.

The reason why plants accumulated such a diversity of different actin isoforms is mysterious. This is exacerbated by the fact that actin sequences are highly conserved and only very few residues mutated to give rise to the different actin isoforms. What could be the biochemical effects of such subtle mutations? We hypothesize that one possible reason for the diversification of actins is to generate cytoskeletal networks with tailored interactomes. Moreover, the mutations have possibly enabled the tailoring of actin-actin interactions in the formation of actin filaments; could different actin isoforms form independent f-actin filaments by suppressing co-polymerization of multiple isoactins into a single filament? There is some evidence for the latter: for AtACT2 and AtACT7 (92.5% homologous) it was proposed that these form independent cytoskeletal structures in *Arabidopsis* cells^29,30^. To explore this notion, we visualised what parts of the actin protein structure are subject to mutational variation. First, we plotted the maximum (and 2^nd^ maximum) amino acid frequency per position as a proxy for conservation (Fig. 3A, Supp. Table 6). This highlights that the N-terminus, and the residues in the region of 210-280 (in the alignment) are the most variable. Visualizing the degree of variability onto the three-dimensional AlphaFold3 structure of the monomeric form of *Arabidopsis* AtACT7, we find that the region that displays the strongest evolutionary variation is an α-helix in subdomain 4 (Fig. 3B). This region, known as the V-stretch, was already known for its variability^3131^. Moreover, the same helix was identified in animal actins in a very different approach, as it was found to be the only site on the actin protein where a small epitope tag could be genetically inserted without interfering with cellular actin function^32^. Given this observation, it is most likely that this helix does not play a key role in actin-protein interactions but rather has a structural function.

**Figure 3.**
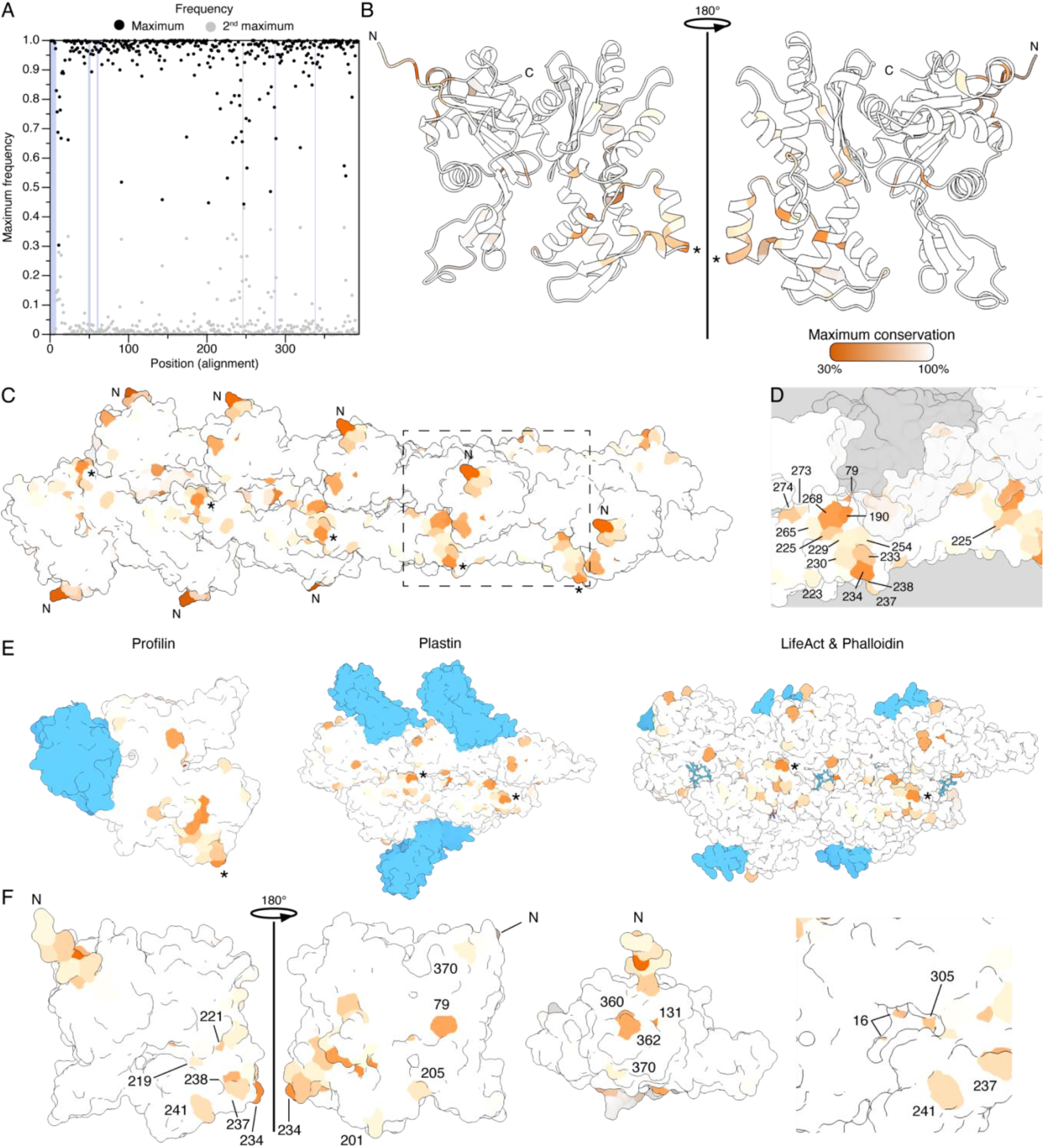
Actin shows distinct variable regions. (A) The percentage of the two most occurring amino acids were plotted over the length of the alignment. The shaded regions (in blue) highlight major gaps in the alignment. (B-E) The conservation rates (as in panel A) plotted on a AlphaFold2 structure of Arabidopsis ACT7 (B) or a crystal structure of filamentous human β-actin (C-E). (E) Structures of major actin-bound actin interactors in blue. Asterisks indicate residue 246 as reference to panels B-D. Note that the highly variable actin N-terminus is missing in the crystal structures of panel E. (F) The variable regions on the surface of the AlphaFold2 structure of Arabidopsis ACT7. Major variable positions are highlighted. Positions 16 and 305 are in the nucleotide binding cleft.

As f-actin is the cytoskeletal form of actin, we also projected the evolutionary variability on the crystal structure of human β-actin (Fig. 3C). Strikingly, we now see that multiple variable positions, that do not connect in the linear sequence (*e*.*g*. position 190, 225 and 265, 268), are in fact in close spatial proximity in the folded f-actin structure (Fig. 3D). In other words, the most variable residues fold together to form a structural zone of maximum mutational variation on the surface of the f-actin filament. This region is not in or near the actin-actin binding interface of f-actin. Hence, this variable region is either causing the specialization of the actin isoforms through their interactomes, or it is not important for actin function and therefore not subject to strong evolutionary pressure. To distinguish these two scenarios, we visualised f-actin-bound crystal structures of two key actin-binding proteins that are conserved across the eukaryotic domain: profilin and fimbrin (known as plastin in metazoa). In addition, we also visualized the well-known actin-binding peptides LifeACT and phalloidin, which are established to bind actin in all eukaryotic lineages (Fig. 3E). None of these 4 interactors bind to the region of maximum variability; nor is this region known as a binding site for other actin-binding proteins. Thus, it is likely that this site is not important for protein-protein interactions. We thus conclude that the variable region might not have any critical function, hence not be subject to strong conservation pressure, and as a result be allowed to mutate relatively free of penalty.

Yet, it is unlikely that actin isoform diversity (Fig. 1A) and its correlation to the complexity of the plant body plan (Fig.1C) occurred without a distinct function. We thus focused on the other, minor, variable positions on the surface (Fig. 3F). Some of these are conserved substitutions between threonine and serine (16, 79, 205, 305); the cumulative frequencies of these 2 amino acids at these positions are close to 100%. Such conserved substitutions are likely to have subtle or no effects on actin polymerisation or protein-protein interactions. This has been studied for position 16, which is in the hydrophobic pocket and essential for binding to ATP. The S16A mutation (S14A in yeast) caused severe heat sensitivity, indicating the importance of this position, but the conserved S16T substitution did not cause any phenotype^33^. Other positions show non-conservative substitutions, like position 79 (52% S, 32% T, 13% N). Interestingly, this position was previously identified as a differentiating epitope to distinguish Type I (+ACT11) and II actins, which contain an S and N on this position, respectively^34^. It is also, among 15 other AAs, a major binding location for phalloidin^35^. However, the biological relevance of this substitution in Type II actins remains unknown. Other positions were more promising for explaining isoform specialization. First, actin polymerisation rates can be reduced by Serine O-GlcNAcylation of S201^36,37^. Serine is the most dominant AA on this location (82%), but a 3 subclusters of the Type I actins contain a Methionine accounting for 12% of the investigated plant actins. This position is adjacent to the T203-205 cluster, which can be phosphorylated to inhibit binding of the Villin/Gelsolin/Fragmin superfamily, and is thus important for f-actin stability^38^. Out of this family, plants only express villins, as actin bundling proteins^39,40^. Within this T203-205 cluster, position 205 is also variable and contains mostly threonines (77%) or serines (22%). Serine phosphorylation of position 241 is believed to stabilize the f-actin structure^41^. Serine is indeed dominant in 73% of the studied plant actins, but 16% of plant actins contains an asparagine in this position. Taken together, some of the variable positions we identify are likely regulators of actin dynamics and/or interactors and are therefore good candidates to understand the functional diversification of actin. It is worth noting, that not every variable position in our analysis is described in literature. The majority of studies into the effects of point mutations in actin come from the medical field and have been identified as the cause of some human pathology. The residues we have identified as being variable in our study of plant actins and which have not been described in literature are thus likely not causing pathological or functional defects, but may in fact be relevant for the functional specification of plant isoactins.

### Actin evolution in the brassicales

As we observe that extant plant actin isoforms diversified and specialized relatively recently, we argue that important substitutions are not easily identified when averaging over all plant actins. Namely, specializations needed in one taxonomic clade might be very different from another clade and averaging over a too large section of the plant lineage obscures these differences. However, comparing actins within a single organism would not allow for the identification of long-term substitutions, but rather non-causal mutations segregating within populations. Hence, to dive further into the possible functions of actin isoform diversification, we decided on an intermediate-scale approach. Our aim is to identify key substitutions by comparing Type I and Type II actins at the eudicot level. Type I and II actins form distinct clades and are not functionally interchangeable^11,12^. In *Arabidopsis thaliana*, mis-expressing a Type II actin under a Type I actin promoter leads to severe actin bundling and growth defects^11,12^. Thus, despite the subtle difference, they are not functionally redundant. To identify possible causative differences, we first investigated the conserved differences between eudicot type I and II actins, selected as those residues that display a |>60%| change in amino acid frequency between Type I and II. This revealed that only seven amino acids are dominantly different. In particular, S360A and G362A caught our attention, as they are close together and are in the extremely conserved C-terminus (Fig. 4A). This region is essential for filamentous actin formation and cannot be modified without losing actin function^32^. Actin undergoes ‘aging’ in its filamentous form: actin monomers in a filament are first ATP bound, followed by an ADP+Pi bound intermediate state and their attachment to the filament becomes unstable after the release of Pi, following by the release of actin monomers^42^. It is this continuous temporal cycle that cause f-actin filaments to treadmill^43^. Actin in the ADP-Pi bound state undergoes a cascade of small structural changes, in which C376 becomes folded into a hydrophobic pocket (formed by Y135, I359, V372). Then, C376 unfolds after the release of Pi and the actin becomes destabilized. Residues 360 and 362 are both polar in Type I actins and hydrophobic in Type II actins. We find that the polar amino acids in Type I actins give rise to an increased number of hydrogen bonds (Fig. 4B). This likely stabilizes this region, and we expect that this affects the hydrophobic pocket that is required for the folding of C376 and may thus alter the treadmilling dynamics Type I vs Type II f-actin.

**Figure 4.**
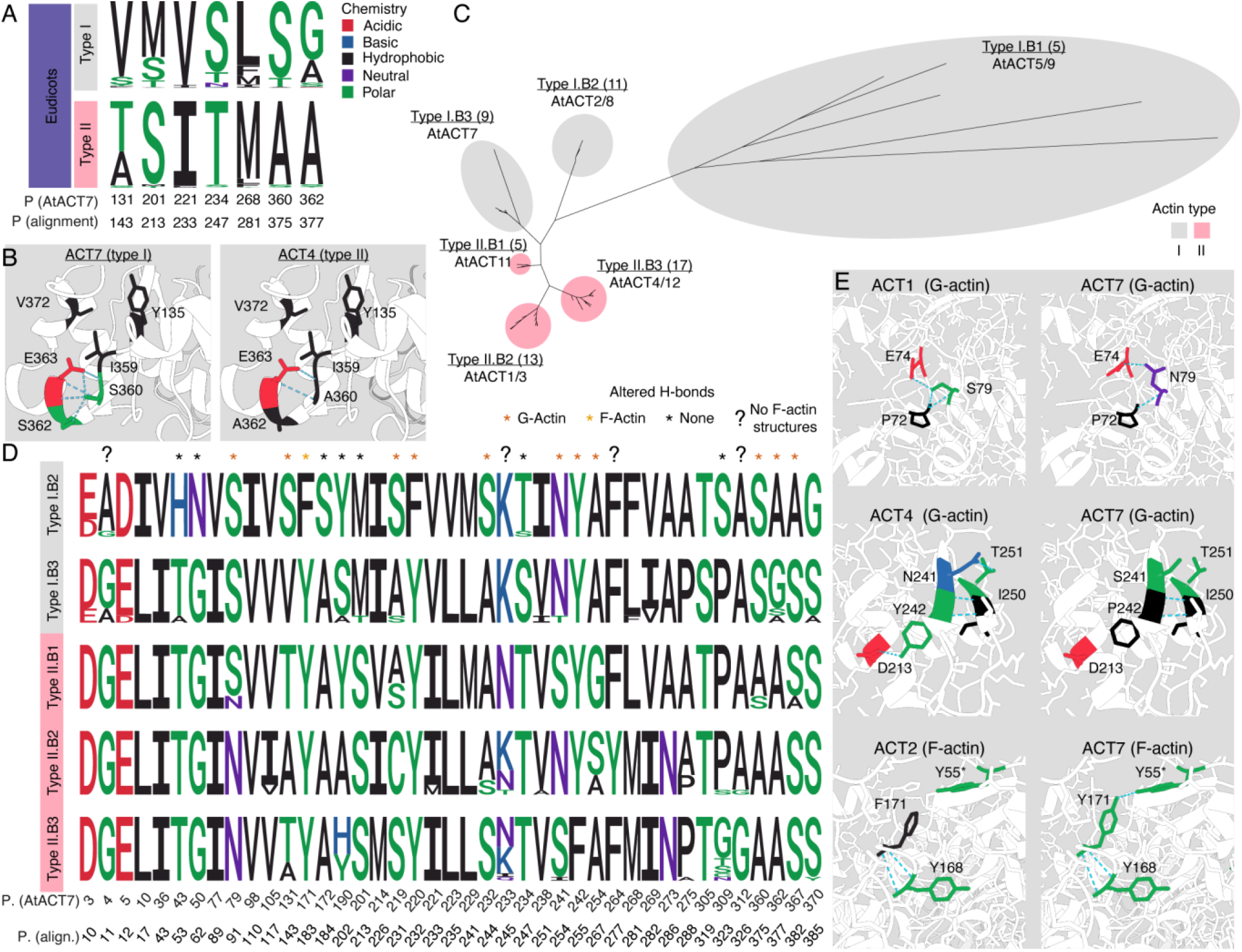
Actin specialization happened after the diversification of the dicots. (A) Major differences between dicot type I and II actins. (B) Unrooted tree for 60 actins of six species in the order brassicales. Arabidopsis actins are written for reference. Number between brackets indicate the number of unique protein sequences. (C) Major amino acid substitutions between the clusters of brassicales. Asterisks indicate differences in predicted hydrogen bonds. (D) AlphaFold2 structures of proteins with differences in predicted hydrogen bonds. Colors indicate the amino acid chemistry.

Type I and II actins are well separated in our phylogenetic tree, which allows for a broader eudicot-wide analysis. Based on our phylogenetic tree (Fig. 1A), we concluded that the divergence of extant actin isoforms and their functional specialization also occurred relatively recently in evolution. For example, we could not detect any other malvids (to which the brassicales belong) near the brassicales cluster. This indicates that the brassicales actins specialized approximately 60 MYA. Hence, we opted for a more fine-grained analysis and considered the 6 species in the brassicales order included in our larger phylogeny. From our overall phylogeny, it is immediately clear that the brassicales actins are all closely related, emphasizing that actin specialized recently in evolution (Supp. Fig. 5). Interestingly, *Arabidopsis* AtACT2 and AtACT8 cluster together with the pseudoactins AtACT5 and AtACT9, showing that these are *not* canonical actins. This is in agreement with functional studies: while the *act7* mutant shows depolymerized actin networks and severe overall growth defects, an *act2/8* double mutant has an almost wild-type like actin structure and whole-plant phenotype, but cannot develop root hairs^14,44–46^.

We first created a phylogenetic tree with the actin sequences of brassicales to identify the actin clusters within this order (Fig. 4C). Type I.B1 has long branches and contains ACT5 and 9, which are likely pseudogenes and were excluded from further analysis. Besides these pseudogenes, five other clusters were identified, which is in line with our earlier phylogeny. Next, we highlighted the major differences (|>60%| change in amino acid ratio) between the clades. Due to this fine-grained approach, we now identified 41 amino acids that are different between these distinct clusters (Fig. 4D). Many, but not all, are conservative substitutions. For now, we focused on the positions which are either polar or hydrophobic and identified 11 substitutions with altered predicted hydrogen bond conformations within the monomer and one (Y171) between two actin subunits in filamentous actin (Fig. 4D+E, Supp. Fig. 6). Y171 (Y169 in humans, F169 in yeast) has been extensively investigated and is essential for actin polymerization due to interactions in the D-loop^47,48^. The mutant F169C cannot polymerize in the absence of phalloidin^48^. Type I.B2 actins are thus more yeast-like and cannot form a hydrogen bond with the Y55. Such hydrogen bonds in a critical region for actin polymerization may cause changes to the actin turnover dynamics and/or the mechanical stability of the filaments. Similar effects are to be expected when position 55 contains a hydrophobic AA. Interestingly, both Y55 and P55 are naturally occurring, and their polymerization dynamics are indeed slightly different^49^. Also, positions 218, and possibly 217, are important in actin polymerization dynamics, due to their effect on the neighboring β-sheet^50^ which is part of the pointed end and essential for intra-strand interactions in f-actin. It has been found that an unstable β-sheet could lead to weaker interactions. Hydrogen bonds might thus stabilize the region and create more stable actin filaments and networks. Taken together, Type I.B2 actins may thus have reduced stability in both the D-loop (position 55), pointy end (position 220) and the C-terminus (position 362). Therefore, we hypothesize that Type I.B2 actins may thus form finer and more transient filaments. This has the conceptual consequence that fine and temporally unstable filaments are not easily visualized with the methods currently available and may thus have remained unobserved^44^. This could explain the observation that Arabidopsis *act2/8* (Type I.B2) mutants still appear to form wildtype-like filaments, while no filaments were found in *act7-4* (Type I.B13) mutants. To the best of our knowledge, not much has been reported on the other amino acids and the effect of small substitutions on actin structure. Taken together, we show that studying actin isoforms at the order level reduces random variation observed when studying a single species, while it may still reveal the essential substitutions that cause functional specialization. Those are not identified when averaging actins on the class or phylum level. Furthermore, we show that small AA substitutions are likely causing subtle changes in the actin dynamics, which may drive actin specialization.

### Conclusions

In this study, we revisited the evolutionary history of plant actins by leveraging the rapidly expanding availability of high-quality genomes across the green lineage. By combining sensitive sequence identification, phylogenetic reconstruction, comparative sequence analysis, and structural mapping, we provide a comprehensive updated view of actin diversification. Our analyses reveal that plant actin isoforms emerged repeatedly and relatively recently in evolutionary history, despite the ancient origin and extreme conservation of the actin fold. We show that actin diversification correlates with key evolutionary transitions. Importantly, we find that only a limited number of conserved substitutions underlie these diversification events, and that these map to specific regions of the actin structure.

A central insight from our work is that actin evolution reflects a balance between stringent conservation of core structural and functional features and localized flexibility that permits diversification. We identify a prominent surface patch that appears largely free from strong evolutionary constraint, suggesting that not all sequence variation is functionally consequential. This is consistent with recent work in animal and yeast actins, in which the same patch was found to be particularly permissible to the introduction of transgenic sequences for isoform-specific actin tagging^32^. By contrast, more subtle and spatially clustered mutational hotspots are enriched at positions likely to influence filament dynamics, stability, and turnover. This supports the notion that isoactin specialization may be driven primarily by tuning intrinsic biophysical properties of f-actin, rather than by large-scale rewiring of the actin interactome. At the same time, our findings do not exclude contributions from isoform-specific protein–protein interactions, and instead suggest that both mechanisms may act in concert.

While our evolutionary and structural analyses generate clear hypotheses, they also highlight a major gap in the field: the lack of direct experimental frameworks to link specific sequence differences to functional outcomes. Addressing this gap will require systematic, isoform-resolved approaches, for which the basis has recently been laid^32^, to reveal whether distinct isoactins engage unique subsets of actin-binding proteins or differ quantitatively in their binding affinities, thereby defining specialized cytoskeletal networks. In addition, given the relatively limited number of amino acid mutations between functionally distinct plant isoactins, it is feasible to pinpoint the causative mutations that lead to functional specialization with targeted mutagenesis, enabling mapping of individual amino acid substitutions to specific biophysical phenotypes. A beautiful example of this can be found in the animal field, where flight-deficient Drosophila mutants in a flight-muscle-specific actin were complemented by site-directed mutated cytosolic actin. A single amino acid substitution in cytosolic actin could largely rescue the flight phenotype, pinpointing a single-residue mutation as a major contributor to the functional nonequivalence of these actin isoforms. Ultimately, integrating isoform-specific interactomics with quantitative biophysics and evolutionary analysis offers a possible route to resolving the long-standing mystery of how subtle changes in one of the most conserved proteins in eukaryotes give rise to the remarkable functional complexity observed in plant cytoskeletons.

## Experimental Methods

### Selection of the proteomes/genomes

First, we selected and downloaded proteomes from major plant lineages. We especially tried to select proteomes to make a balanced representation of the diversity in the major plants lineages. This was an iterative process, starting with e.g. all 123 proteomes from Bryogenomes^51^. However, such approach did not resolve the actin evolution better due to the high redundancy in genera. Hence, we decided to allow one proteome per genus and ideally less than 4 per family. Exceptions were made based on the importance of the family. We included 15 Poaceae (of 15 different genera) as their type II actins form a specific cluster and more species were required for our analyses. Six brassicales (of 6 genera) were added because Arabidopsis has the best described actins. We made a special effort to include lycophytes, ferns and gymnosperms as only few genomes are described and these form major transitions in plant evolution. Actins of two lycophytes (*Lycopodium clavatum* and *Isoetes taiwanensis*) were extracted from the genomes and we used annotated proteomes for all other species. We relied on various resources like Uniprot (59 proteomes), Bryogenomes (38), JGI (36), PLAZA (27), Fernbase (6), Hornwortbase (6) and others (7) (Supp. Table 1).

### Protein sequence extraction and alignment

Protein sequences were extracted using a custom made python pipeline. First, five actins (*Arabidopsis thaliana* ACT2, *Marchantia polymorpha* ACT1, *Phytophthora infestans* ACTa, *Homo sapiens* ACTB and *Saccharomyces cerevisiae* ACT) were aligned using Clustal Omega and used to create a Hidden Markov Model (HMM) profile. Due to the high conservation of actin sequences, this minimalistic set was sufficient to detect highly divergent actins of Chlamydomonas and reported pseudogenes (e.g. AtACT5 and 9). Hence, we argued that adding more actin sequences to the profile did not improve the HMM search. Then the proteomes described in the previous section were scanned using this profile and hits were filtered for a full sequence hit score > 70. Next, we used the filtered hits for an HMM search using different existing profiles for actin (PTHR11937-SF572), ARP1 (PTHR11937-SF155), ARP2 (PTHR11937-SF37:SF37), ARP3 (PTHR11937-SF31), ARP4 (PTHR11937-SF413), ARP6 (PTHR11937-SF47), ARP7 (PTHR11937-SF46), ARP9 (PTHR11937-SF574), β-centractin (PTHR22629). The HMM scores were, together with the sequence length, used for PCA analysis, which was clustered using FactoMiner^52^. Group 6 contained actins and 2 human centractins, which were removed based on the HMM score for β-centractin (score >800).

Next, we started with the curation of Group 6 actin sequences. First, we flagged all sequences with a C-terminus other than “CF” or with a length outside the range of [370-380] AAs and aligned them individually to Arabidopsis ACT2, which was used as reference. We argued that the start codon was misannotated if the flagged sequence (1) had an extension at the N-terminus and (2) a methionine was aligned to the start codon of AtACT2. Alternatively, we argued that the stop codon was misannotated if the flagged sequence had (1) an extension at the C-terminus and (2) and the sequence contained the canonical actin C-terminus (HRKCF) that was aligned to the AtACT2 reference. Sequences with N- or C-terminus extensions that matched these criteria were truncated to match the AtACT2 reference. We also removed presumably falsely annotated introns when the alignment with AtACT2 showed gaps with a length > 5 amino acids. A maximum of 1 intron was removed per sequence. Next, we flagged the sequences that still not matched the criteria and used the genomic data of the organism (NCBI tBLASTn, v2.17.0+) to identify the correct protein sequence. Lastly, the sequences that still not matched the criteria were manually reconstructed using downloaded genomic data. 171 actin sequences were curated using these methods and 42 were impossible. Next, we aligned the sequences with ClustalOmega (v1.2.4). We noticed that especially the N-terminus was very gappy, which we restored by moving all gaps in the first 15 amino acids (*e*.*g*. ------MS----LS to MSLS). Other misaligned gaps were aligned to the AtACT2 reference. We argue that the curation of gaps is important for such conserved proteins because, unlike less conserved proteins, these gaps contain important biological information. The phylogenetic tree was created with IQ-TREE2 (v2.2.0)^53^ with 1000 bootstraps. The tree was rooted on the *Candidatus Lokiarchaeum ossiferum* actin 2951803_UPI0027A42375. We first used ModelFinder to identify the best model, and validated two models: the top-ranked model according to the Corrected Akaike Information Criterion (LG+G4) and Bayesian Information Criterion (Q.insect+I+I+R7, Supp. Table 7). Our most important findings (as summarized in Fig. 1B) were supported by both models, however the Q.insect+I+I+R7 model resulted in some unexpected results (Supp. Fig. 7). Most importantly, the Brassicales cluster Type I.B2 (containing AtACT2 and 8, Fig. 4C) was placed among the type II actins, which does not correspond to existing literature^17,19^. In addition two actins clustered with type II with LG+G4 and type I with Q.insect+I+I+R7. Sequence analysis (A360/S360) revealed that those are likely angiosperm type IIs (A360). Therefore, we decided to use the LG+G4 model. The phylogenetic tree with Brassicales actins was also created with this model.

### LOGO and ChimeraX

Amino acid frequencies were calculated with the Biostrings (v2.66.0)^54^ PSSM function and visualized with the ggseqlogo package (v0.2)^55^. Evolutionary conserved substitutions were identified by comparing all lineages that formed before- and after the divergence of a new lineage, and was iterated for all major lineages. For example, to identify the conserved substitution in the lycophytes: (prasinodermophyta-bryophyte) were grouped and compared with those in lineages that formed after this formation (lycophytes-eudicot). Type I and II actins were grouped separately.

The g-ACT visualizations were made with the AtACT7 AlphaFold2 structure (AF-A0A178UEW4-F1) using ChimeraX-1.9^56^. Human ACTB (3LUE^57^) was used for the visualizations on f-act. Hydrogen bonds were predicted in AlphaFold2 models of AtACT2 (AF-Q96292-F1), 7 (AF-P53492-F1), 11 (AF-P53496-F1), 1 (AF-P0CJ46-F1) and 4 (AF-P53494-F1) using ChimeraX. Hydrogen bonds in the f-actin structure were predicted in the SWISS-MODELs of ACT2 (Q96292_7-377:8trm.1.C) and ACT7 (P53492_7-377:8trm.1.C). These were also experimentally validated before^47^.

## Data availability

Processed trees can be found in https://itol.embl.de/shared/JasperLamers

## Acknowledgements

This work and J.L. and J.S. are funded by the European Research Council, project Catch, project number 101000981.

---

**Supplemental Figure 1.**
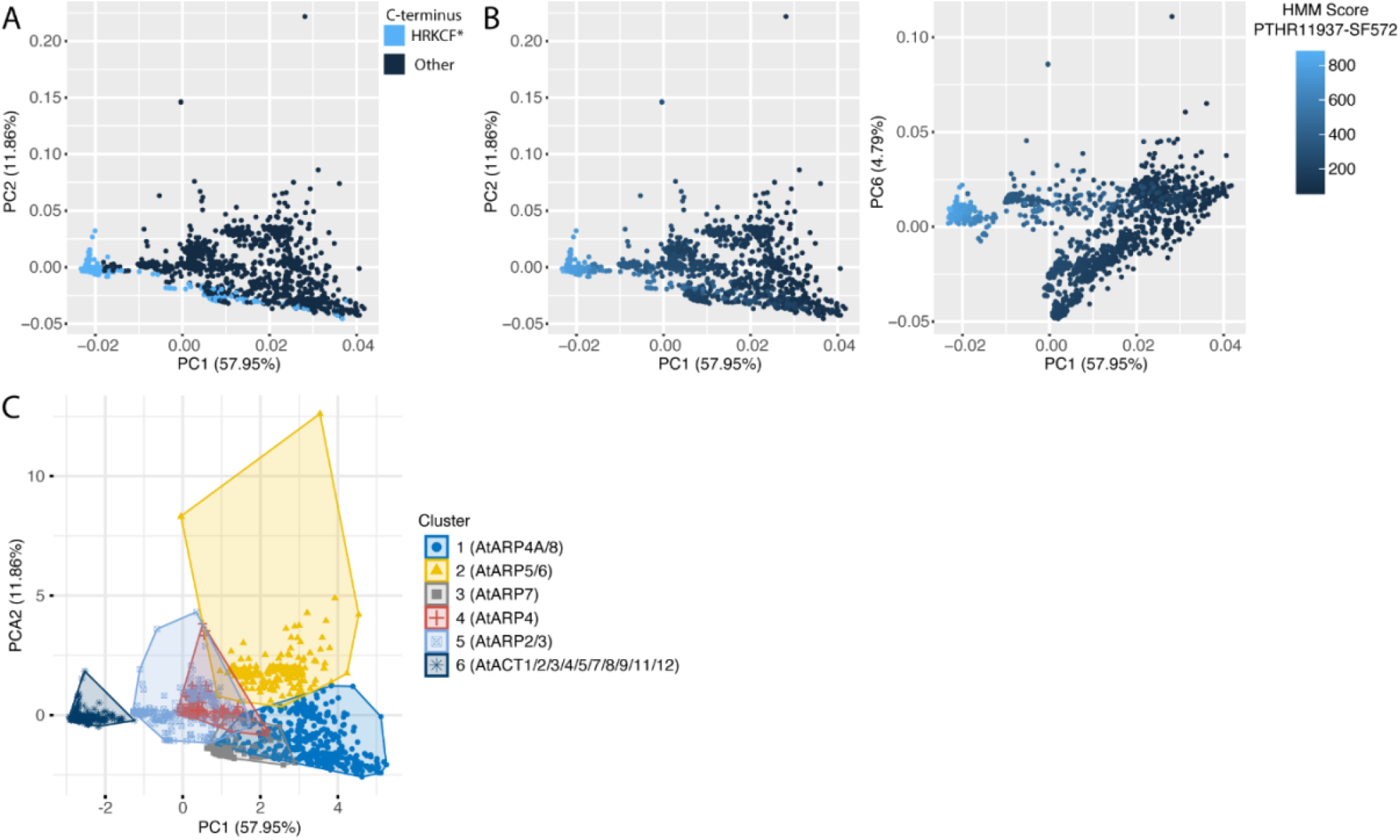
Actins were separated from actin related proteins (ARPs) using a PCA plot of the HMM scores. (A-C) PCA plots of the HMM scores of existing panther profiles for actin, ARP1, ARP2, ARP3, ARP4, ARP6, ARP9 and β-centractin. Colorcoding proteins with the conserved actin C-terminus (HRKCF). The C-terminus of plant ARP7 is equal to actin, which are the scattered points. (A) or the HMM scores for actin (B). (C) Clusters were identified in the PCA plot and cluster 6 contains the proteins with the conserved C-terminus and the highest HMM scores for actin.

**Supplemental Figure 2.**
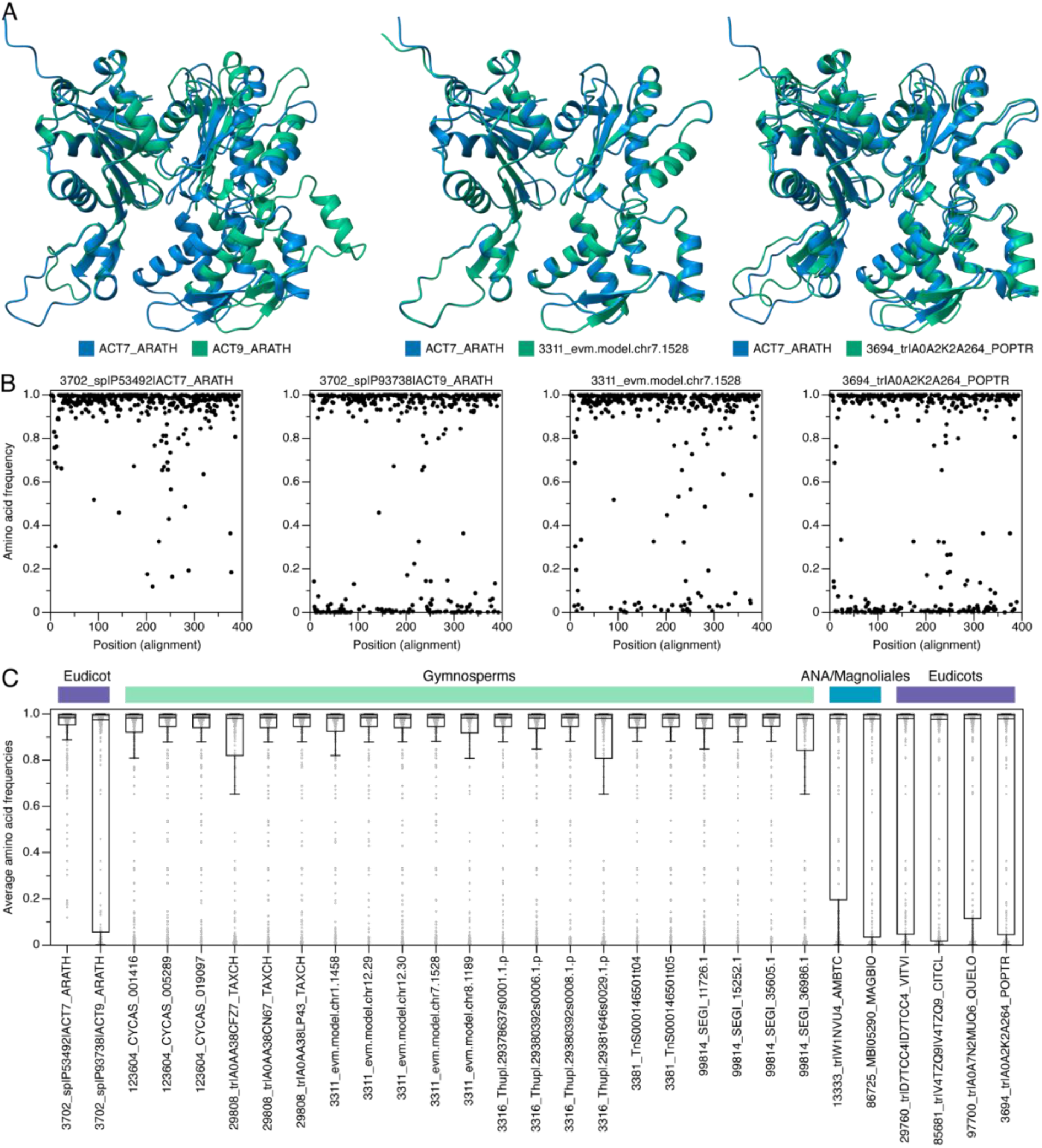
The gymnosperm actins in cluster I.G are real actins. In our phylogeny we discovered a deviant cluster of actins, which contained gymnosperm and a few dicot actins. (A) We first randomly picked a gymnosperm (3311_evm.model.chr7.1528) and a dicot (3694_tr|A02K2A264_POPTR) actin from this cluster and predicted their structure with AlphaFold3. Those were compared to AtACT7, which is known to be a functional actin. AtACT9 is described as a pseudogene and was used to validate our methodology. While the gymnosperm actin nearly equals AtACT7, the dicot and AtACT9 actins both showed structural differences. (B-C) Next, we compared the sequences of these actins. For each alignment position (including gaps), we calculated how often this AA was identified at that position across all other investigated plant actins. While the gymnosperm actin showed some unusual AAs, AtACT9 and 3694_tr|A02K2A264_POPTR showed many of such locations (B). We extended this calculation to all actins in Cluster I.G and visualized this in a boxplot. This reveals that the dicot actins generally contain more unusual AAs and are comparable to the ATACT9 pseudogene.

**Supplemental Figure 3.**
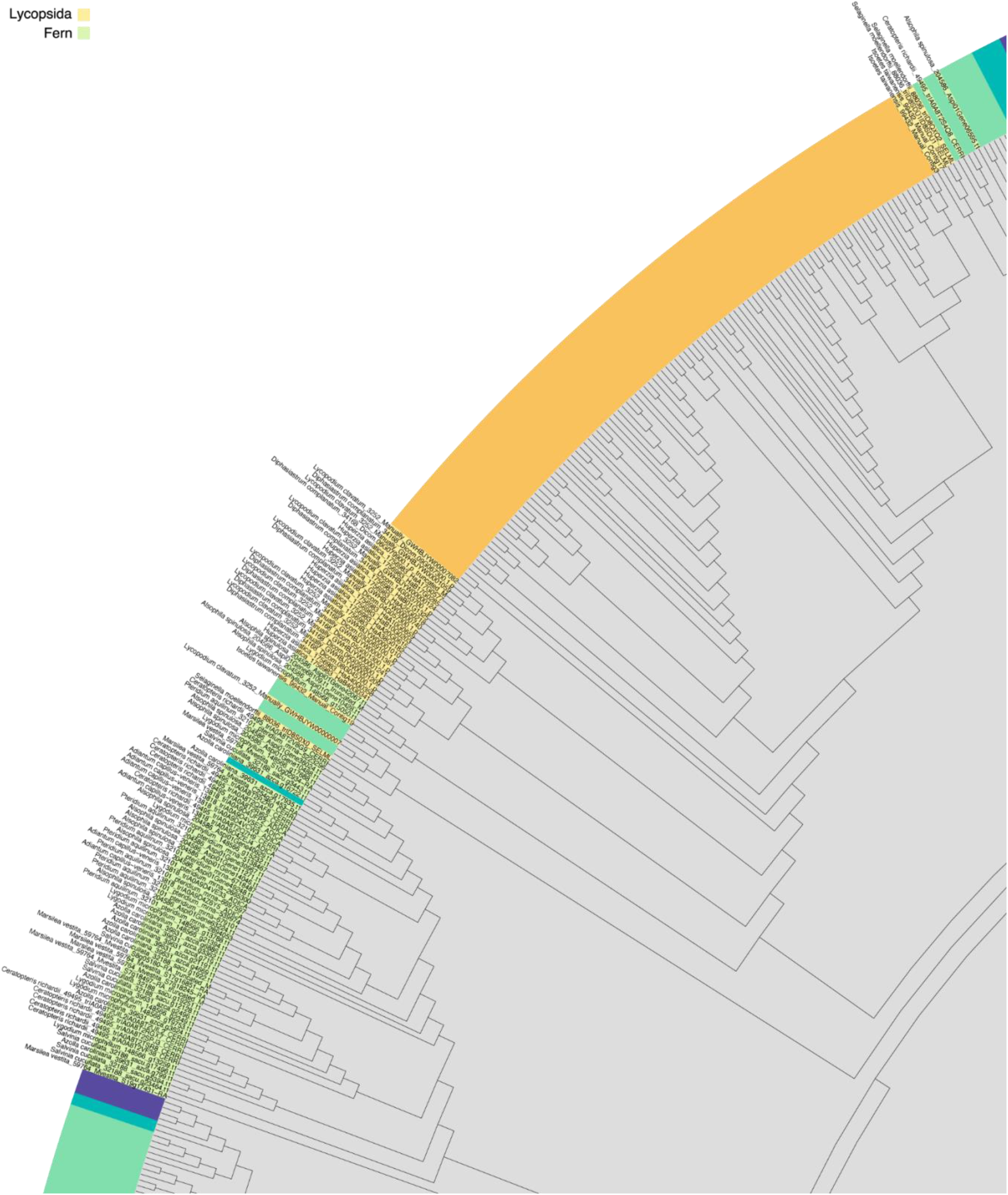
Lycophyte and fern actins have often duplicated, but not diverged. Here the lycohpytes and ferns are highlighted in zoom of the phylogenetic tree shown in Fig. 1A.

**Supplemental Figure 4.**
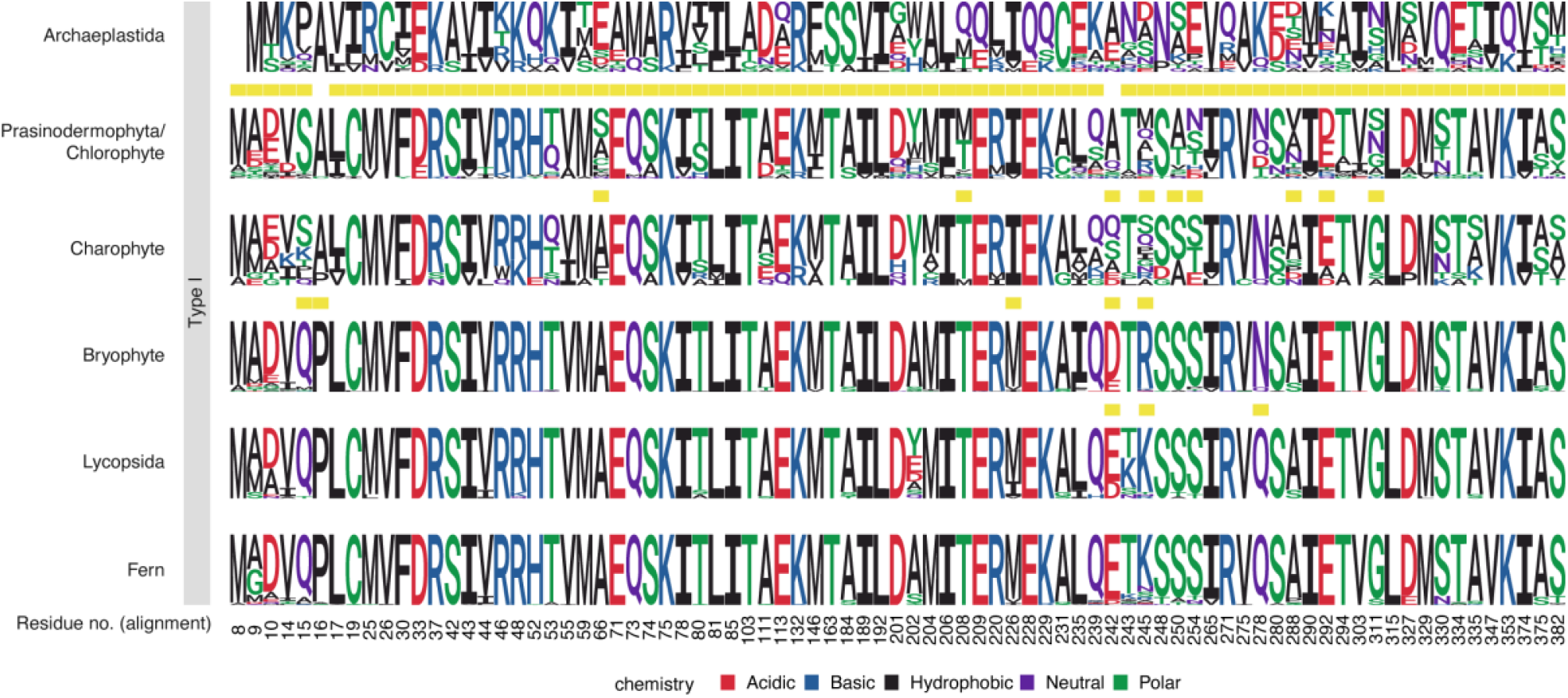
Major conserved transitions in the Type I actin sequences are mapped per major taxonomic clade. In contrast to Figure 2, the Archaeaplastida are included in the comparison.

**Supplemental Figure 5.**
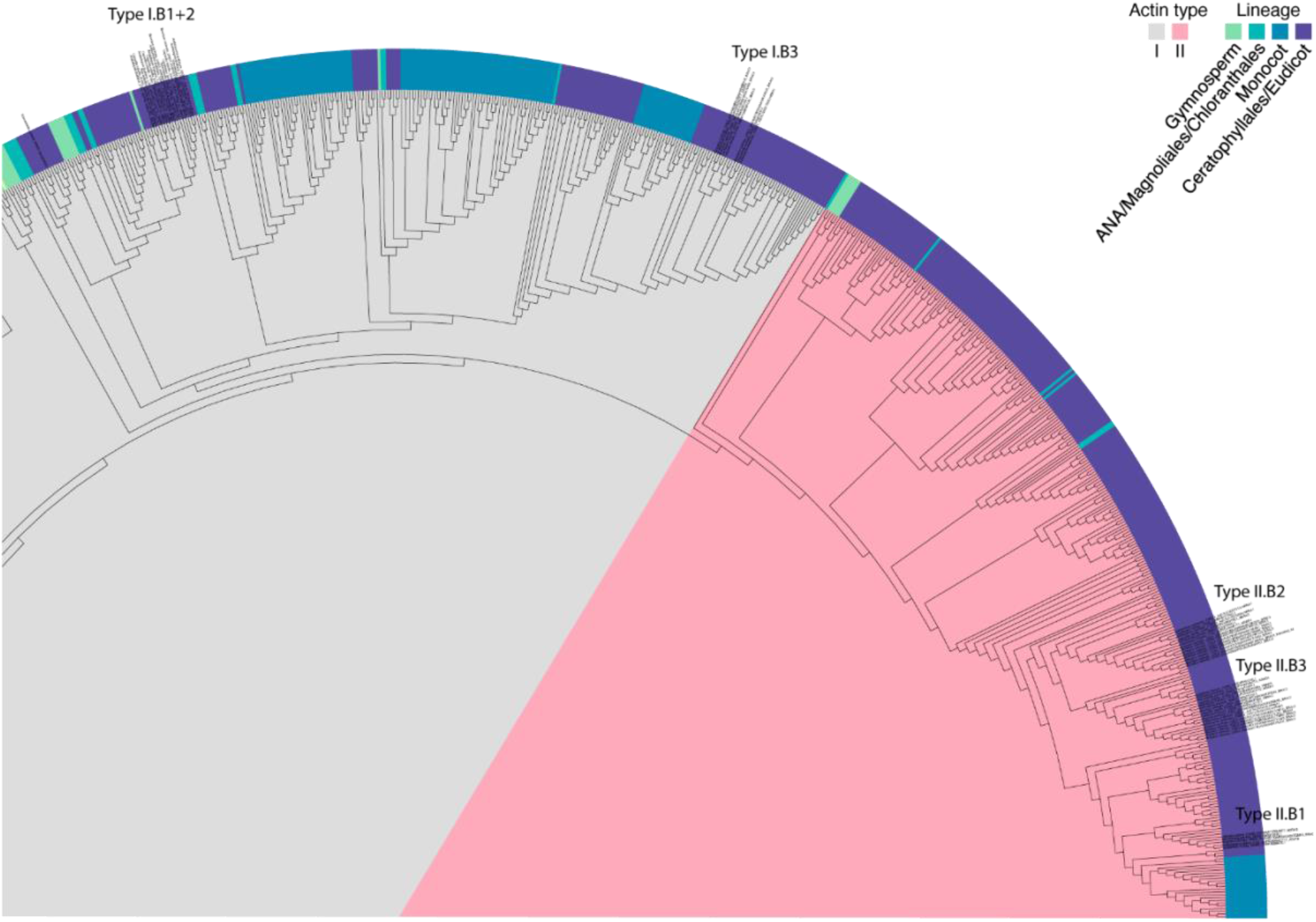
Eudicot actins have diverged recently in evolution. Here the actins in the brassicales order are highlighted in zoom of the phylogenetic tree shown in Fig. 1A.

**Supplemental Figure 6.**
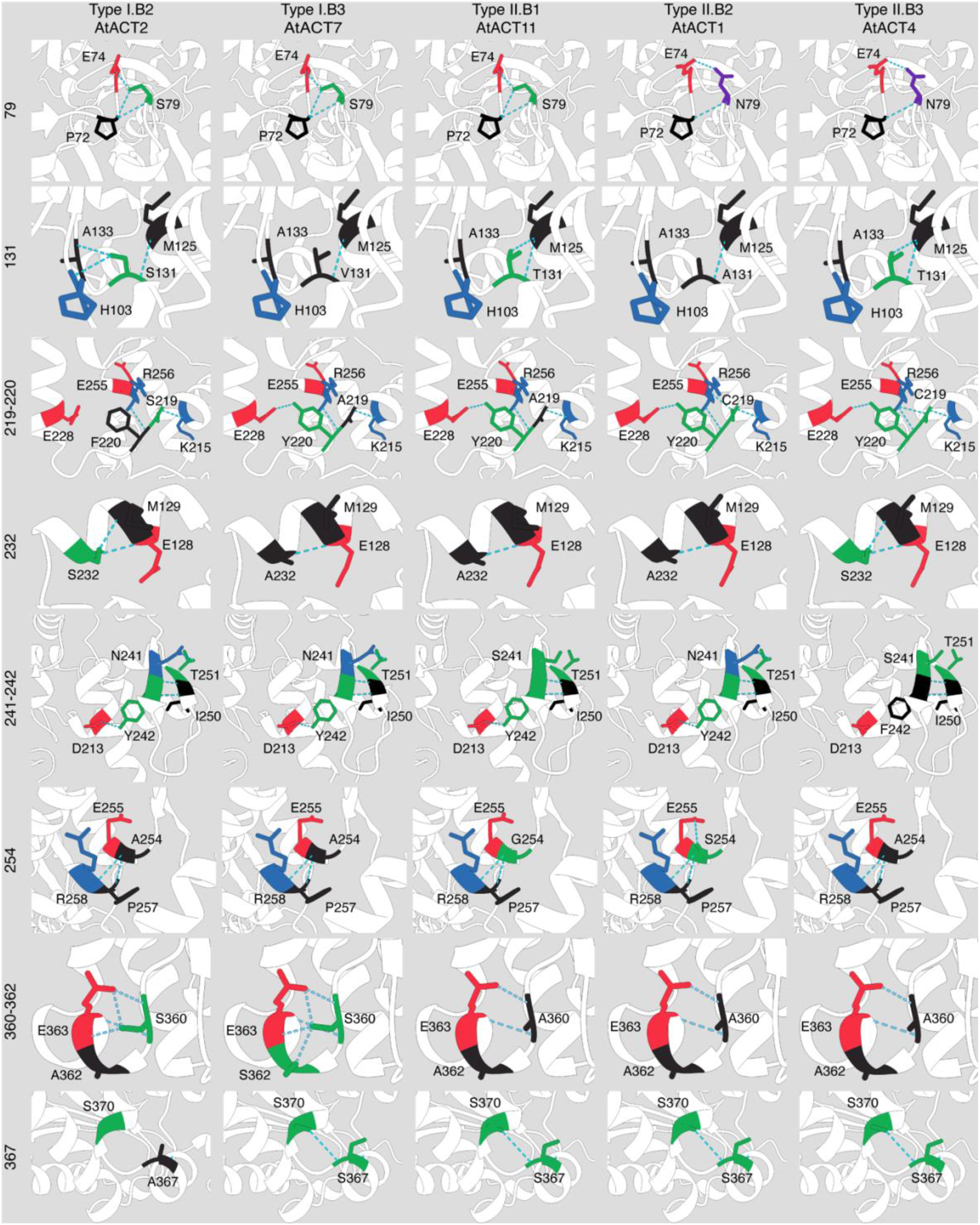
Brassicales actin clusters have distinct hydrogen bond conformations. All substitutions that affect hydrogenbond conformations (Fig. 4D) are shown here. One Arabidopsis actin was choosen per cluster as defined in Fig. 4C.

**Supplemental Figure 7.**
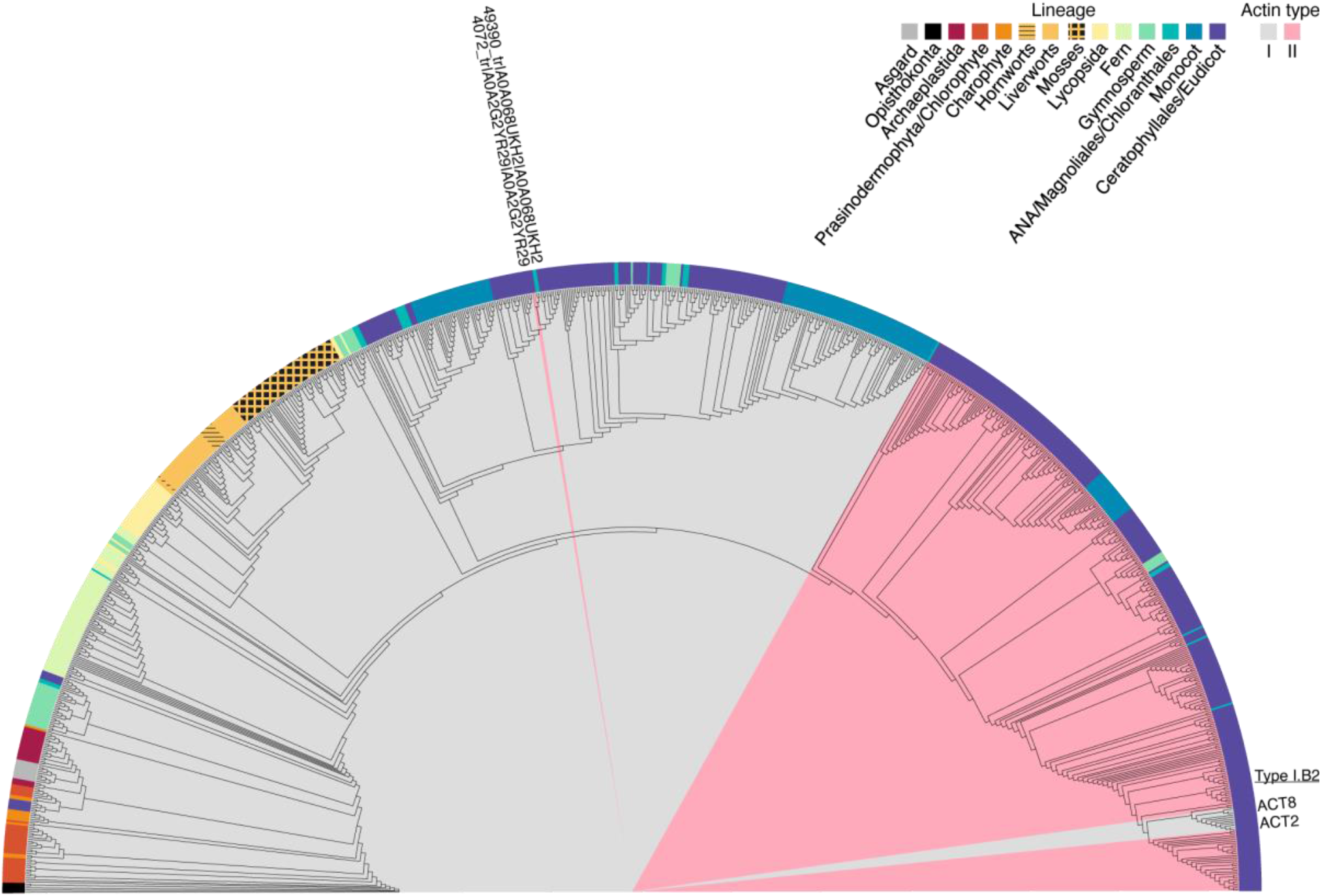
The choice of model used in IQ-TREE2 does not affect major findings. This phylogenetic tree was made using the best model according to the Bayesian Information Criterion (Q.Insect+I+I+7). We used the type annotation of Fig. 1A to highlight the major differences between these models. Our most important findings (e.g. segregation per plant lineage, type II actins in the gymnosperms, cluster I.G and that actins diversified and specialized recently) are supported by both models. The major difference is that the Brassicales cluster Type I.B2 is now grouped with the type II actins. This is not supported by literature^17,19^. Minorly, two type II actins are now grouped with type I actins, both actins contain A360, which is canonical for angiosperm type II actins (Fig. 2).

## References

1. Rodrigues-Oliveira, T. et al. Actin cytoskeleton and complex cell architecture in an Asgard archaeon. Nature 613, 332–339 (2023).

2. An, Y. et al. Strong, constitutive expression of the Arabidopsis ACT2/ACT8 actin subclass in vegetative tissues. Plant J. 10, 107–121 (1996).

3. Huang, S., An, Y.-Q., McDowell, J. M., McKinney, E. C. & Meagher, R. B. The Arabidopsis ACT11 actin gene is strongly expressed in tissues of the emerging inflorescence, pollen, and developing ovules. Plant Mol. Biol. 33, 125–139 (1997).

4. Šlajcherová, K., Fišerová, J., Fischer, L. & Schwarzerová, K. Multiple actin isotypes in plants: Diverse genes for diverse roles? Front. Plant Sci. 3, (2012).

5. Vandekerckhove, J., Bugaisky, G. & Buckingham, M. Simultaneous expression of skeletal muscle and heart actin proteins in various striated muscle tissues and cells. A quantitative determination of the two actin isoforms. J. Biol. Chem. 261, 1838–1843 (1986).

6. Nowak, K. J. et al. Rescue of skeletal muscle alpha-actin-null mice by cardiac (fetal) alpha-actin. J. Cell Biol. 185, 903–915 (2009).

7. Jaeger, M. A., Sonnemann, K. J., Fitzsimons, D. P., Prins, K. W. & Ervasti, J. M. Context-dependent functional substitution of alpha-skeletal actin by gamma-cytoplasmic actin. FASEB J. Off. Publ. Fed. Am. Soc. Exp. Biol. 23, 2205–2214 (2009).

8. Kumar, A. et al. Rescue of cardiac alpha-actin-deficient mice by enteric smooth muscle gamma-actin. Proc. Natl. Acad. Sci. U. S. A. 94, 4406–4411 (1997).

9. Patrinostro, X. et al. Essential nucleotide- and protein-dependent functions of Actb/β-actin. Proc. Natl. Acad. Sci. U. S. A. 115, 7973–7978 (2018).

10. Vedula, P. et al. Diverse functions of homologous actin isoforms are defined by their nucleotide, rather than their amino acid sequence. eLife 6, e31661 (2017).

11. Kandasamy, M. K., McKinney, E. C. & Meagher, R. B. Functional Nonequivalency of Actin Isovariants in Arabidopsis. Mol. Biol. Cell 13, 251–261 (2002).

12. Meagher, R. B., Kandasamy, M. K. & McKinney, E. C. Multicellular development and protein-protein interactions. Plant Signal. Behav. 3, 333–336 (2008).

13. Kandasamy, M. K., McKinney, E. C. & Meagher, R. B. A single vegetative actin isovariant overexpressed under the control of multiple regulatory sequences is sufficient for normal Arabidopsis development. Plant Cell 21, 701–718 (2009).

14. Kandasamy, M. K., McKinney, E. C., Roy, E. & Meagher, R. B. Plant vegetative and animal cytoplasmic actins share functional competence for spatial development with protists. Plant Cell 24, 2041–2057 (2012).

15. Ringli, C., Baumberger, N. & Keller, B. The Arabidopsis Root Hair Mutants der2–der9 are Affected at Different Stages of Root Hair Development. Plant Cell Physiol. 46, 1046–1053 (2005).

16. The Arabidopsis Genome Initiative. Analysis of the genome sequence of the flowering plant Arabidopsis thaliana. Nature 408, 796–815 (2000).

17. An, S. S., Möpps, B., Weber, K. & Bhattacharya, D. The Origin and Evolution of Green Algal and Plant Actins. Mol Biol Evol 16, 275–285 (1999).

18. The Arabidopsis Genome Initiative. Analysis of the genome sequence of the flowering plant Arabidopsis thaliana. Nature 408, 796–815 (2000).

19. Mcdowell, J. M., Huang, S., Mckinney, E. C.An, Y.-Q. & Meaghed, R. B. Structure and Evolution of the Actin Gene Family in Arabidopsis Thalianu. https://academic.oup.com/genetics/article/142/2/587/6016673 (1996).

20. Gerrienne, P., Meyer-Berthaud, B., Fairon-Demaret, M., Street, M. & Steemans, P. Runcaria, a middle devonian seed plant precursor. Science 306, (2004).

21. Meagher, R. B., McKinney, E. C. & Vitale, A. V. The evolution of new structures: clues from plant cytoskeletal genes. Trends Genet. 15, 278–284 (1999).

22. Cvrčková, F., Bezvoda, R. & Žárský, V. Computational identification of root hair-specific genes in Arabidopsis. Plant Signal. Behav. 5, 1407–1418 (2010).

23. Abramson, J. et al. Accurate structure prediction of biomolecular interactions with AlphaFold 3. Nature 630, 493–500 (2024).

24. Hightower, R. C. & Meagher, R. B. THE MOLECULAR EVOLUTION OF ACTIN. Genetics 114, 315–332 (1986).

25. Mounier, N., Gouy, M., Mouchiroud, D. & Prudhomme, J. C. Insect muscle actins differ distinctly from invertebrate and vertebrate cytoplasmic actins. J. Mol. Evol. 34, 406–415 (1992).

26. Onishi, M., Pringle, J. R. & Cross, F. Evidence that an unconventional actin can provide essential F-actin function and that a surveillance system monitors F-actin integrity in Chlamydomonas. Genetics 202, 977–996 (2016).

27. Jack, B., Mueller, D. M., Fee, A. C., Tetlow, A. L. & Avasthi, P. Partially Redundant Actin Genes in Chlamydomonas Control Transition Zone Organization and Flagellum-Directed Traffic. Cell Rep. 27, 2459-2467.e3 (2019).

28. Szoevenyi, P. Genome assemblies and annotations of the three Anthoceros accessions as well as alignment matrices and tree files used for reconstructing the land plant phylogeny. 161088987 Bytes figshare 10.6084/M9.FIGSHARE.9974999 (2020).

29. Kijima, S. T., Hirose, K., Kong, S. G., Wada, M. & Uyeda, T. Q. P. Distinct biochemical properties of Arabidopsis thaliana actin isoforms. Plant Cell Physiol. 57, 46–56 (2016).

30. Kijima, S. T. et al. Arabidopsis vegetative actin isoforms, AtACT2 and AtACT7, generate distinct filament arrays in living plant cells. Sci. Rep. 8, (2018).

31. Splettstoesser, T., Holmes, K. C., Noé, F. & Smith, J. C. Structural modeling and molecular dynamics simulation of the actin filament. Proteins Struct. Funct. Bioinforma. 79, 2033–2043 (2011).

32. van Zwam, M. C. et al. IntAct: A nondisruptive internal tagging strategy to study the organization and function of actin isoforms. PLoS Biol. 22, (2024).

33. Chen, X. & Rubenstein, P. A. A Mutation in an ATP-binding Loop of Saccharomyces cerevisiae Actin (S14A) Causes a Temperature-sensitive Phenotype in Vivo and in Vitro. J. Biol. Chem. 270, 11406–11414 (1995).

34. Kandasamy, M. K., McKinney, E. C. & Meagher, R. B. The late pollen-specific actins in angiosperms. Plant J. 18, 681–691 (1999).

35. Kumari, A., Kesarwani, S., Javoor, M. G., Vinothkumar, K. R. & Sirajuddin, M. Structural insights into actin filament recognition by commonly used cellular actin markers. EMBO J. 39, e104006 (2020).

36. Akimoto, Y. et al. The O-GlcNAcylation of β-actin Ser199 controls nuclear speckle localization and is dysregulated in diabetes. Histochem. Cell Biol. 163, 106 (2025).

37. Akimoto, Y. et al. O -GlcNAcylation and phosphorylation of β-actin Ser^199^ in diabetic nephropathy. Am. J. Physiol.-Ren. Physiol. 317, F1359–F1374 (2019).

38. Furuhashi, K., Hatano, S., Ando, S., Nishizawa, K. & Inagaki, M. Phosphorylation by actin kinase of the pointed end domain on the actin molecule. J. Biol. Chem. 267, 9326–9330 (1992).

39. Klahre, U., Friederich, E., Kost, B., Louvard, D. & Chua, N.-H. Villin-Like Actin-Binding Proteins Are Expressed Ubiquitously in Arabidopsis. Plant Physiol. 122, 35–48 (2000).

40. Van Der Honing, H. S., Kieft, H., Emons, A. M. C. & Ketelaar, T. Arabidopsis VILLIN2 and VILLIN3 Are Required for the Generation of Thick Actin Filament Bundles and for Directional Organ Growth. Plant Physiol. 158, 1426–1438 (2012).

41. El-Khobar, K. E. et al. Polo-like kinase-1 mediates hepatitis C virus-induced cell migration, a drug target for liver cancer. Life Sci. Alliance 6, e202201630 (2023).

42. Oosterheert, W., Klink, B. U., Belyy, A., Pospich, S. & Raunser, S. Structural basis of actin filament assembly and aging. Nature 611, 374–379 (2022).

43. Neuhaus, J.-M., Wanger, M., Keiser, T. & Wegner, A. Treadmilling of actin. J. Muscle Res. Cell Motil. 4, 507–527 (1983).

44. Numata, T., Sugita, K., Ahamed Rahman, A. & Rahman, A. Actin isovariant ACT7 controls root meristem development in Arabidopsis through modulating auxin and ethylene responses. J. Exp. Bot. 73, 6255–6271 (2022).

45. Ringli, C., Baumberger, N., Diet, A., Frey, B. & Keller, B. ACTIN2 Is Essential for Bulge Site Selection and Tip Growth during Root Hair Development of Arabidopsis. Plant Physiol. 129, 1464–1472 (2002).

46. Nishimura, T., Yokota, E., Wada, T., Shimmen, T. & Okada, K. An Arabidopsis ACT2 Dominant-Negative Mutation, which Disturbs F-actin Polymerization, Reveals its Distinctive Function in Root Development. Plant Cell Physiol. 44, 1131–1140 (2003).

47. Hvorecny, K. L., Sladewski, T. E., De La Cruz, E.M., Kollman, J. M. & Heaslip, A. T. Toxoplasma gondii actin filaments are tuned for rapid disassembly and turnover. Nat. Commun. 15, (2024).

48. Kudryashov, D. S., Grintsevich, E. E., Rubenstein, P. A. & Reisler, E. A nucleotide state-sensing region on actin. J. Biol. Chem. 285, 25591–25601 (2010).

49. Liu, X., Shu, S., Hong, M.-S. S., Yu, B. & Korn, E. D. Mutation of Actin Tyr-53 Alters the Conformations of the DNase I-binding Loop and the Nucleotide-binding Cleft. J. Biol. Chem. 285, 9729–9739 (2010).

50. Kotila, T. et al. Structural basis of rapid actin dynamics in the evolutionarily divergent Leishmania parasite. Nat. Commun. 13, 3442 (2022).

51. Dong, S. et al. Bryophytes hold a larger gene family space than vascular plants. Nat. Genet. 57, 2562–2569 (2025).

52. Lê, S., Josse, J. & Husson, F. FactoMineR : An R Package for Multivariate Analysis. J. Stat. Softw. 25, (2008).

53. Minh, B. Q. et al. IQ-TREE 2: New Models and Efficient Methods for Phylogenetic Inference in the Genomic Era. Mol. Biol. Evol. 37, 1530–1534 (2020).

54. H. Pagès, P. A. Biostrings. Bioconductor 10.18129/B9.BIOC.BIOSTRINGS (2017).

55. Wagih, O. ggseqlogo: A ‘ggplot2’ Extension for Drawing Publication-Ready Sequence Logos. 0.2.2 10.32614/CRAN.package.ggseqlogo (2017).

56. Goddard, T. D. et al. UCSF ChimeraX: Meeting modern challenges in visualization and analysis. Protein Sci. 27, 14–25 (2018).

57. Galkin, V. E., Orlova, A., Salmazo, A., Djinovic-Carugo, K. & Egelman, E. H. Opening of tandem calponin homology domains regulates their affinity for F-actin. Nat. Struct. Mol. Biol. 17, 614–616 (2010).

